# Conserved statistical organization of cis-regulatory landscapes defines a fluctuation-structured regulatory phase

**DOI:** 10.64898/2026.02.06.704399

**Authors:** Yoshihiro Ohmori

## Abstract

Cis-regulatory elements undergo extensive evolutionary turnover, particularly among enhancers, yet gene expression programs remain remarkably conserved across species. This paradox suggests that regulatory conservation may reside not in individual DNA elements but in higher-order organizational principles of regulatory sequence space. Here, we analyze gene-centered cis-regulatory islands (GCICs) across rice and *Arabidopsis* genomes to characterize the macroscopic statistical organization of cis-regulatory landscapes. GCICs serve as gene-centered representations of combinatorial motif-family overlap that resolve intrinsic sequence organization rather than discrete functional elements. Across both genomes, combinatorial vocabulary diversity scales sublinearly with motif-family abundance, motif-family combinations exhibit conserved hierarchical rank–frequency organization, dominant combinatorial identities are extensively shared with near-proportional quantitative correspondence, and higher-order GCIC architectures composed of multiple islands per gene are stably maintained. Disruption of native sequence structure through randomization induces systematic collapse of higher-order GCIC architectures and drives motif-family combinatorial interactions toward independence, indicating that regulatory landscape organization emerges from intrinsic hierarchical sequence correlations. Together, these conserved scaling laws, hierarchical distributions, stable combinatorial identities, and characteristic responses to perturbation define a conserved statistical regulatory phase underlying cis-regulatory organization across plant genomes. This macroscopic regulatory architecture provides a probabilistic substrate for redundant regulatory potential, enabling functional cis-regulatory elements to emerge and stabilize despite extensive enhancer turnover. Our findings establish that cis-regulatory conservation operates at the level of conserved statistical landscape organization rather than individual regulatory sequences, offering a quantitative framework for understanding regulatory robustness and evolutionary flexibility.

## Introduction

Gene regulation has traditionally been conceptualized as being encoded by discrete cis-regulatory elements such as enhancers and promoters, whose conserved DNA sequences specify regulatory function. Within this framework, preservation of regulatory output was expected to correlate closely with conservation of regulatory sequences themselves.

Comparative regulatory genomics has substantially revised this view. Extensive gain and loss of enhancer elements—referred to as enhancer turnover—has been documented across diverse animal and plant lineages, even where gene expression programs remain broadly conserved across species (Wittkopp & Kalay, 2011; Villar et al., 2015). This apparent decoupling between sequence-level regulatory turnover and conserved transcriptional output has increasingly been attributed to redundant regulatory architectures and network-level buffering, in which multiple regulatory inputs collectively stabilize gene expression programs (Frankel et al., 2010; Osterwalder et al., 2018).

High-throughput functional genomics has further revealed that regulatory activity is broadly distributed across genomic sequence space and depends strongly on local sequence context and combinatorial motif organization rather than isolated high-affinity binding sites (Arnold et al., 2013; Melnikov et al., 2012; Muerdter et al., 2015). Enhancer-like regulatory landscapes have now been uncovered in plants using STARR-seq and chromatin-based approaches, demonstrating that distributed and distal regulatory activity is a conserved feature across eukaryotes (Sun et al., 2019; Oka et al., 2017; Ricci et al., 2019).

Together, these findings suggest that regulatory conservation may operate not primarily at the level of individual cis-regulatory elements, but at higher-order organizational scales characterized by combinatorial structure and statistical regularities.

Recently, we introduced Gene-centered cis-regulatory islands (GCICs) as a gene-centered framework for identifying combinatorial cis-regulatory landscapes from intrinsic local sequence features (Ohmori, 2026). GCICs capture probabilistic overlap of motif families in gene-centered coordinates, providing a complementary representation to element-centric annotations.

Here, we investigate whether regulatory conservation manifests as conserved macroscopic statistical organization of GCIC landscapes across plant genomes. By comparing rice and *Arabidopsis*, we examine (i) scaling relations between motif-family abundance and combinatorial vocabulary diversity, (ii) hierarchical rank–frequency organization of motif-family combinations, (iii) genomic compartmental deployment of GCICs, (iv) conservation and quantitative correspondence of dominant combinations, and (v) higher-order GCIC architectural organization across genes.

Rather than focusing on individual regulatory elements, this study characterizes conserved statistical regimes of gene-centered regulatory landscapes that emerge from intrinsic sequence organization.

## Methods

### Identification of GCIC landscapes

Gene-centered cis-regulatory islands (GCICs) were generated genome-wide for rice and *Arabidopsis thaliana* using the GCIC detection framework described in Ohmori (2026). For rice, the IRGSP-1.0 reference genome and gene annotation were obtained from Ensembl Plants (release 58; files: Oryza_sativa.IRGSP-1.0.dna_sm.toplevel.fa and Oryza_sativa.IRGSP-1.0.58.gtf) (Kawahara et al., 2013; Howe et al., 2020). For *Arabidopsis*, gene-centered regions were constructed using the TAIR10 reference genome and gene annotation obtained from Ensembl Plants (release 62; files: Arabidopsis_thaliana.TAIR10.dna_sm.toplevel.fa and Arabidopsis_thaliana.TAIR10.62.gtf) (Berardini et al., 2015; Howe et al., 2020).

All GCIC detection procedures followed Ohmori (2026), with species-specific sliding window and step size parameters fixed based on experimentally characterized cis-regulatory loci. For rice, a 200 bp window with 50 bp step size was used to recapitulate regulatory architecture at the *DROOPING LEAF* (*DL*) locus (Ohmori et al., 2011), whereas for Arabidopsis, a 100 bp window with 25 bp step size was used to resolve regulatory structure at the *AGAMOUS* (*AG*) locus (Deyholos & Sieburth, 2000). All cross-genome analyses were performed under these fixed resolutions.

GCIC detection was applied within defined gene-centered 5′–3′ coordinates of annotated genes using only the forward genomic strand, yielding a conservative estimate of cis-regulatory landscape organization. Genome-wide GCIC annotations for rice were previously reported (Ohmori, 2026). For *Arabidopsis*, genomic coordinates and properties of gene-centered regions are provided in Supplementary Data 1, and GCIC island annotations with genomic compartment classifications are provided in Supplementary Data 2.

GCICs were operationally defined following Ohmori (2026) as gene-centered genomic regions identified by the spatial overlap of independently enriched motif-family–specific intervals, and were annotated by motif-family composition and genomic compartment (promoter, exon, intron, or intergenic).

### Vocabulary scaling analysis

To quantify combinatorial landscape expansion, we analyzed the relationship between the total number of motif-family occurrences per gene (N) and the number of unique motif families represented within GCIC islands (V).

For each species, gene-level data were log-binned across the full dynamic range of N using geometrically spaced bins. Within each bin, median V values and interquartile ranges were computed.

Scaling relationships of the form *V* = *aN*^*b*^ were fitted by linear regression in log–log space using binned median values. Scaling exponents (b) and coefficients of determination (R^2^) were estimated from these statistics and used as macroscopic descriptors of combinatorial organization.

### Rank–frequency organization of motif-family combinations

Frequency distributions of motif-family combinations were computed across all GCIC islands for each species. Combination identities were normalized to be order-invariant by sorting motif-family components within each combination prior to counting.

Unique combinations were ranked by decreasing frequency, and normalized frequencies were plotted against rank on logarithmic axes. Hierarchical scaling in the dominant combinatorial regime was quantified by linear regression of log-transformed frequency versus log-transformed rank over ranks ≥10, with upper bounds capped by the total number of unique combinations in each genome.

Slopes of these regressions were reported as hierarchical scaling exponents (α), with goodness-of-fit assessed by R^2^.

### Genomic compartment distribution of GCICs

GCICs were classified into promoter, exon, intron, and intergenic regions using gene annotation frameworks for each species. Fractional compositions across compartments were computed and visualized as normalized distributions to compare regulatory landscape deployment between genomes.

### Cross-genome conservation of combinatorial identities

Motif-family combinations were normalized by component ordering and compared across species. Rank–frequency tables were generated independently for rice and *Arabidopsis*.

Overlap of dominant combinations was quantified using Jaccard similarity indices across increasing rank thresholds (K), with K capped by the minimum number of unique combinations observed between species.

Quantitative correspondence of shared combinations was assessed by linear regression in log–log space between normalized usage frequencies in rice and *Arabidopsis*. Regression slopes and R^2^ values were used to evaluate proportional conservation of combinatorial deployment. Combination-level statistics are provided in Supplementary Table S1.

### Randomization analysis of GCIC architectural organization

Higher-order GCIC architectural organization and internal combinatorial structure were assessed by quantifying GCIC multiplicity per gene and deviations of motif-family combination frequencies from independence in both real genomes and randomized sequence controls.

For each random replicate, gene-centered input sequences were randomized by mononucleotide shuffling, preserving regional length and mononucleotide composition while disrupting higher-order sequence correlations. Randomized sequence controls were generated for rice only using gene-centered input sequences corresponding to a randomly selected subset of 5,000 rice genes (Supplementary Data 3). Motif occurrences were recomputed on the shuffled sequences prior to GCIC reconstruction. Ten independent randomized replicates were generated using fixed pseudorandom seeds. GCIC landscapes from randomized subsets were compared against those derived from the same non-randomized rice gene subset, the full rice genome, and the full *Arabidopsis* genome. GCIC detection was performed using identical parameters for all randomized and non-randomized datasets.

Distributions of GCIC multiplicity were compared across real and randomized datasets in the low-order regime (GCIC = 1–4) and the heavy-tail regime (GCIC ≥ 3). Jensen–Shannon divergence was computed using the full multiplicity distributions to quantify distributional distances between real, randomized, and cross-species landscapes (Lin, 1991).

### Analysis of combinatorial deviations from independence

To quantify internal combinatorial organization within GCIC islands, observed frequencies of motif-family combinations were compared against expectations under an independence model.

For each species and dataset, expected combination frequencies were computed as products of marginal motif-family frequencies. Residuals were defined as:

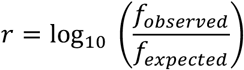

Marginal motif-family frequencies were computed as the fraction of GCIC islands containing each motif family within the same dataset, and expected frequencies for a given combination were calculated as the product of the corresponding marginals.

Dominant motif-family combinations were defined as the top 60 most frequent combinations in the real rice dataset and were used consistently across all datasets for comparative analyses.

Dominant combinations were classified as enriched (positive residuals), depleted (negative residuals), or approximately independent (near-zero residuals), using a threshold of eps = 0.1.

To assess erosion of combinatorial structure under randomization, residuals were computed for each random replicate. Shrinkage toward independence was quantified as the median of |r_random|/|r_real| across dominant motif-family combinations, restricted to those with |r_real| ≥ eps, where r_random and r_real denote residuals in randomized and real datasets, respectively. Combinations absent in randomized datasets were assigned residual values of zero when computing shrinkage, reflecting complete erosion toward independence.

Distributions of shrinkage values across random replicates were visualized using boxplots, and real rice and *Arabidopsis* values were overlaid for comparison. Empirical p-values were computed from the distribution of shrinkage values across randomized replicates.

## Results

GCIC landscapes in rice and *Arabidopsis* display conserved scaling laws of combinatorial expansion, hierarchical rank–frequency organization of motif-family combinations, shared dominant combinatorial identities, and higher-order GCIC architectural structure across multiple quantitative descriptors (Figs. 1–4).

**Figure 1.**
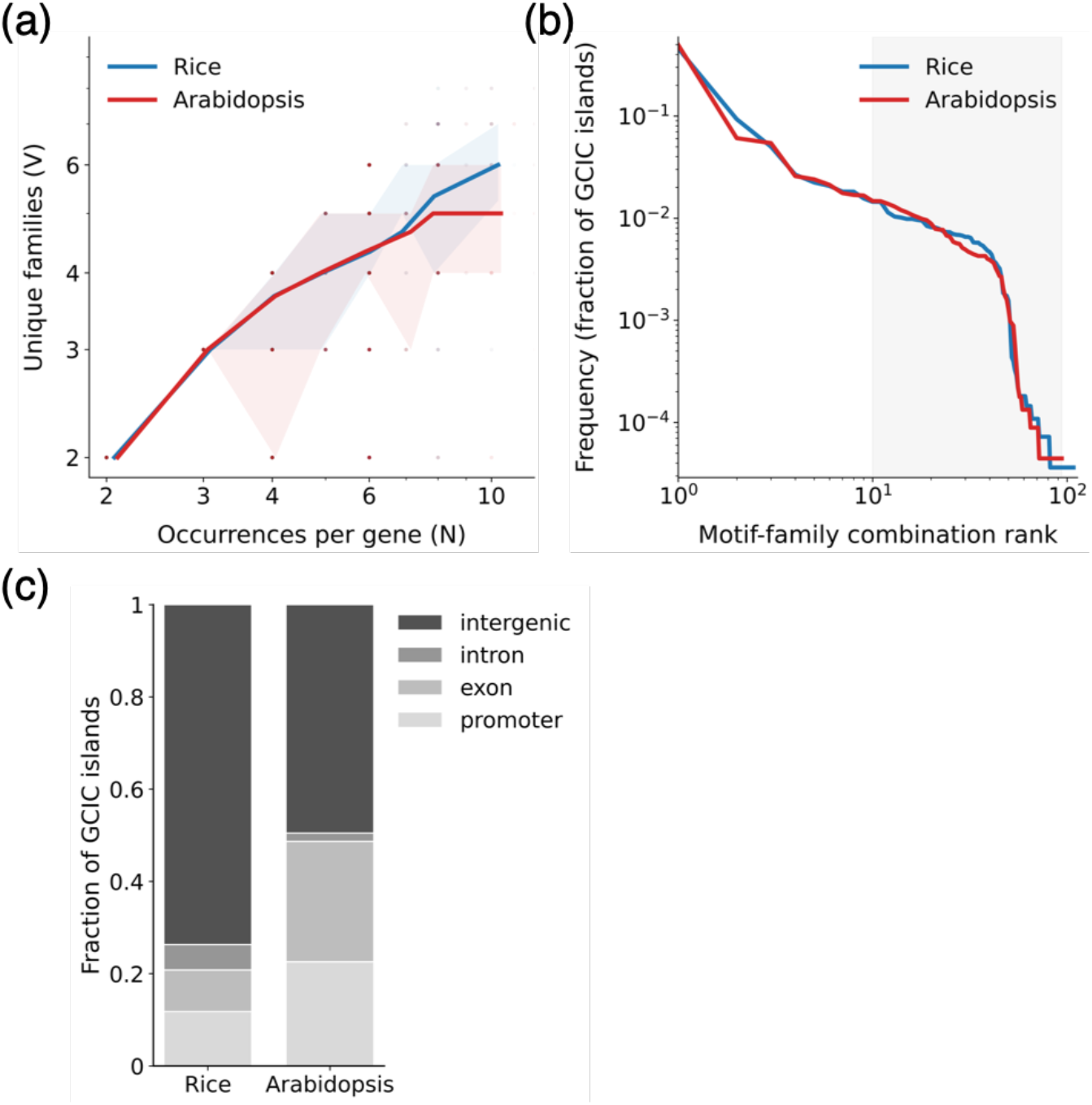
Quantitative organization of gene-centered cis-regulatory island (GCIC) landscapes in rice and *Arabidopsis*. **(a) Vocabulary scaling of GCIC combinatorial landscapes**. Relationship between the total number of motif-family occurrences per gene (N) and the number of unique motif families represented within GCIC islands (V). Gene-level data were log-binned across N, with median V values plotted and interquartile ranges indicated by shaded regions. Solid lines represent fitted sublinear scaling relations of the form *V* = *aN*^*b*^, revealing conserved expansion of combinatorial diversity with increasing regulatory complexity in both genomes. **(b) Hierarchical rank–frequency organization of motif-family combinations**. Normalized frequencies of order-invariant motif-family combinations defining GCIC composition, ranked by decreasing occurrence across all GCIC islands. Distributions span multiple orders of magnitude and exhibit power-law-like scaling in dominant regimes (shaded region), indicating strongly skewed hierarchical usage of combinatorial identities. **(c) Genomic compartment distribution of GCIC islands**. Normalized fractions of GCICs localized within promoter, exon, intron, and intergenic regions for rice and *Arabidopsis*. GCICs are broadly deployed across gene-centered genomic space, with predominant localization outside core promoters and substantial enrichment in intergenic regions.

Scaling relationships between motif-family abundance and combinatorial vocabulary diversity exhibit reproducible sublinear behavior in both genomes across the full dynamic range analyzed (Fig. 1a). Frequency distributions of motif-family combinations follow strongly skewed hierarchical rank–frequency patterns characterized by a small number of dominant combinations and a long tail of rare combinations (Fig. 1b).

GCIC islands are broadly distributed across genomic compartments, with predominant localization outside core promoters and the largest fraction residing in intergenic regions in both genomes (Fig. 1c). A substantial fraction of dominant motif-family combinations is shared between rice and *Arabidopsis* across multiple rank thresholds, and shared combinations display strong quantitative correspondence in usage frequencies across species (Fig. 2a,b).

**Figure 2.**
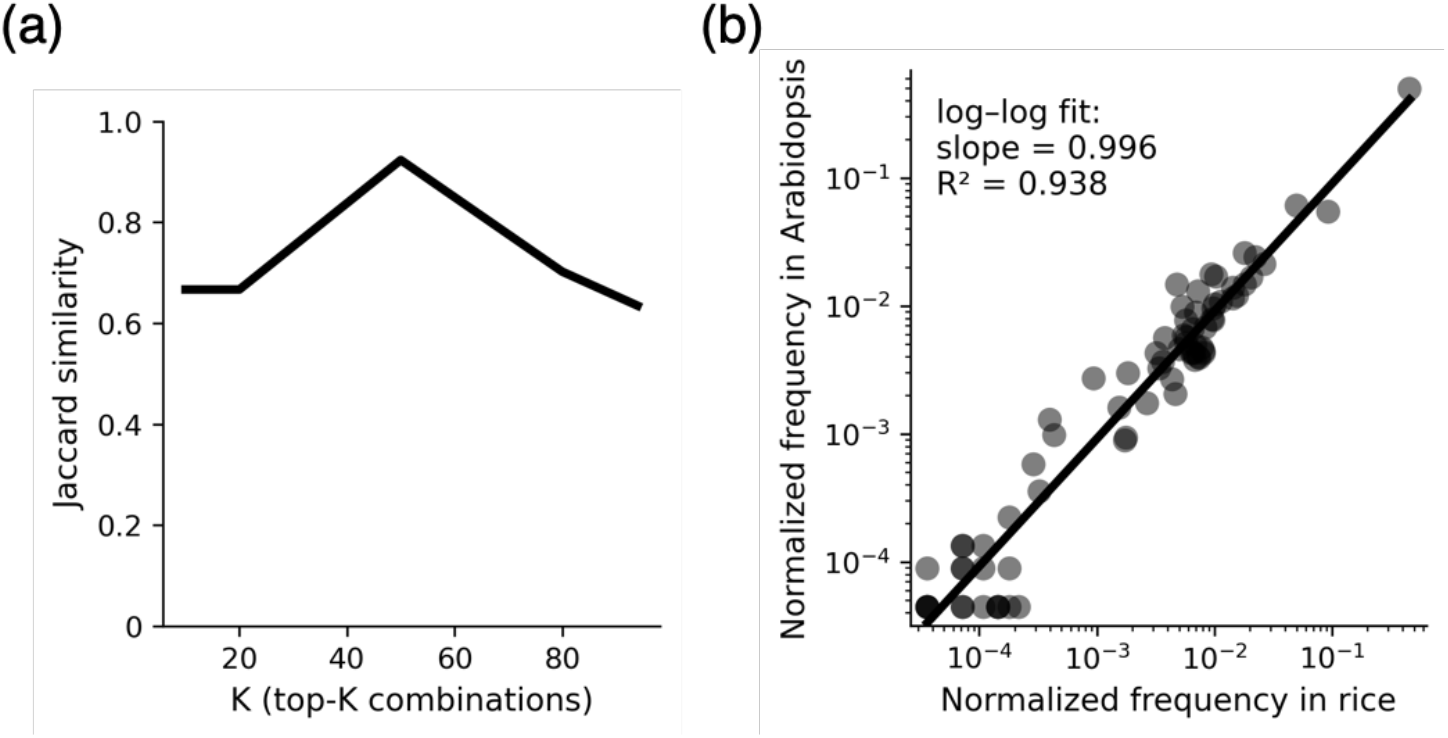
Conservation and proportional correspondence of dominant GCIC combinatorial identities between rice and *Arabidopsis*. **(a) Overlap of dominant motif-family combinations across genomes**. Jaccard similarity of motif-family combination sets between rice and *Arabidopsis* across increasing rank thresholds (K), where combinations were independently ranked by decreasing frequency in each genome. High similarity across broad rank regimes indicates conservation of core combinatorial identities. **(b) Quantitative correspondence of shared combination usage frequencies**. Log–log scatter comparison of normalized frequencies for motif-family combinations shared between rice and *Arabidopsis*. Each point represents a conserved combination, with axes indicating genome-specific frequencies across all GCIC islands. The fitted regression demonstrates near-proportional correspondence across multiple orders of magnitude, indicating quantitative conservation of combinatorial deployment.

Higher-order GCIC architectures composed of multiple islands per gene are stably maintained in real genomes but systematically collapse under sequence randomization (Fig. 3). Internal motif-family combinatorial organization exhibits structured deviations from independence in real genomes and flows toward independence under randomization (Fig. 4a,b).

**Figure 3.**
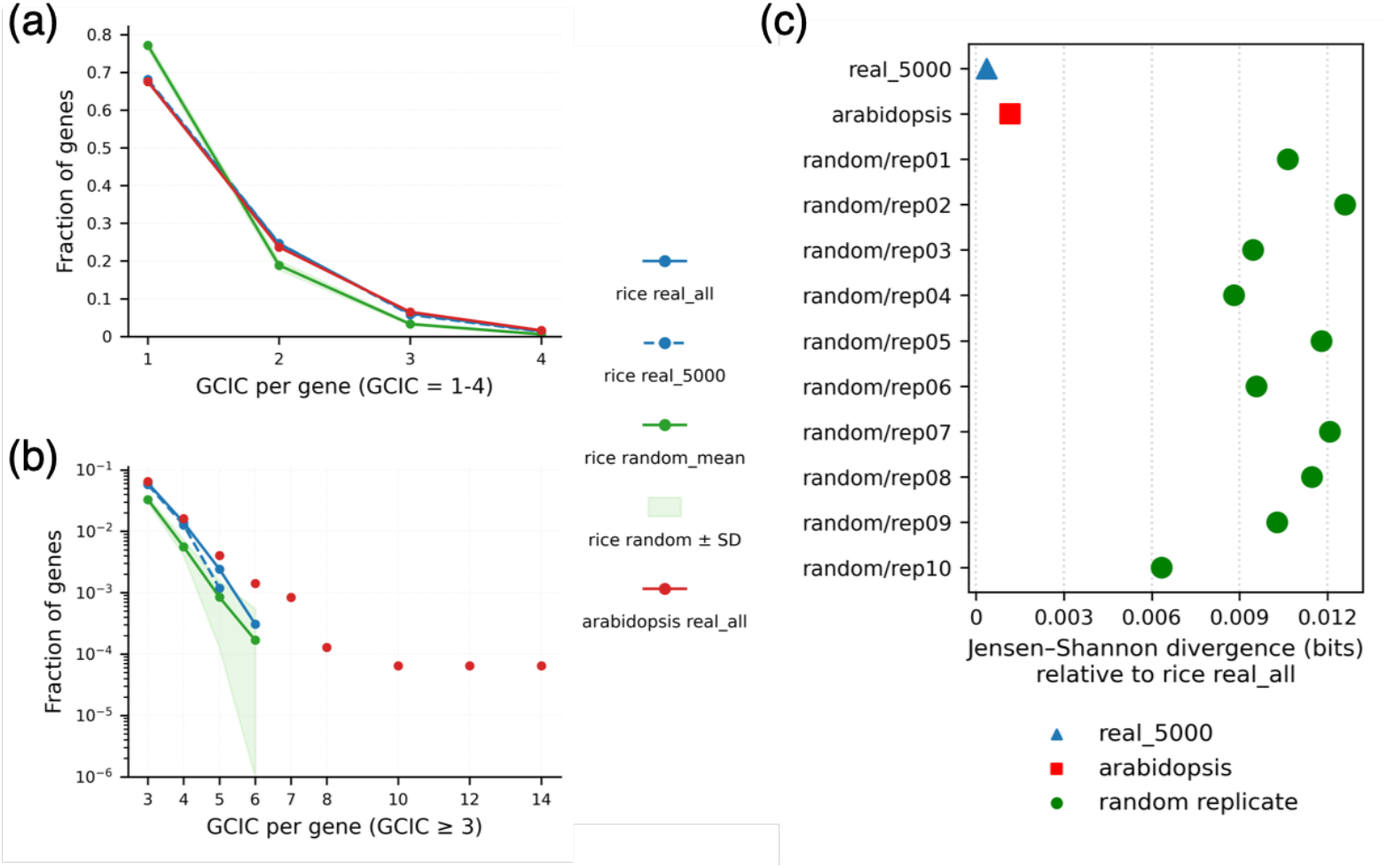
Higher-order GCIC architectural organization collapses under sequence randomization. **(a) Low-order GCIC multiplicity distributions**. Fraction of genes harboring 1–4 GCIC islands in real rice and *Arabidopsis* genomes compared with randomized sequence controls generated from a 5,000-gene rice subset. Real genomes exhibit substantial multi-island architectures, whereas randomization enriches single-island configurations. **(b) Heavy-tail distributions of GCIC multiplicity**. Log-scale distributions of genes containing multiple GCIC islands (GCIC ≥ 3), highlighting higher-order architectural regimes preserved in real genomes but strongly depleted under randomization. **(c) Quantitative divergence of architectural landscapes**. Jensen–Shannon divergence between GCIC multiplicity distributions of real genomes, *Arabidopsis*, and randomized rice replicates relative to real rice landscapes. Randomized controls consistently diverge from real genomic organization, whereas cross-species landscapes remain highly similar.

**Figure 4.**
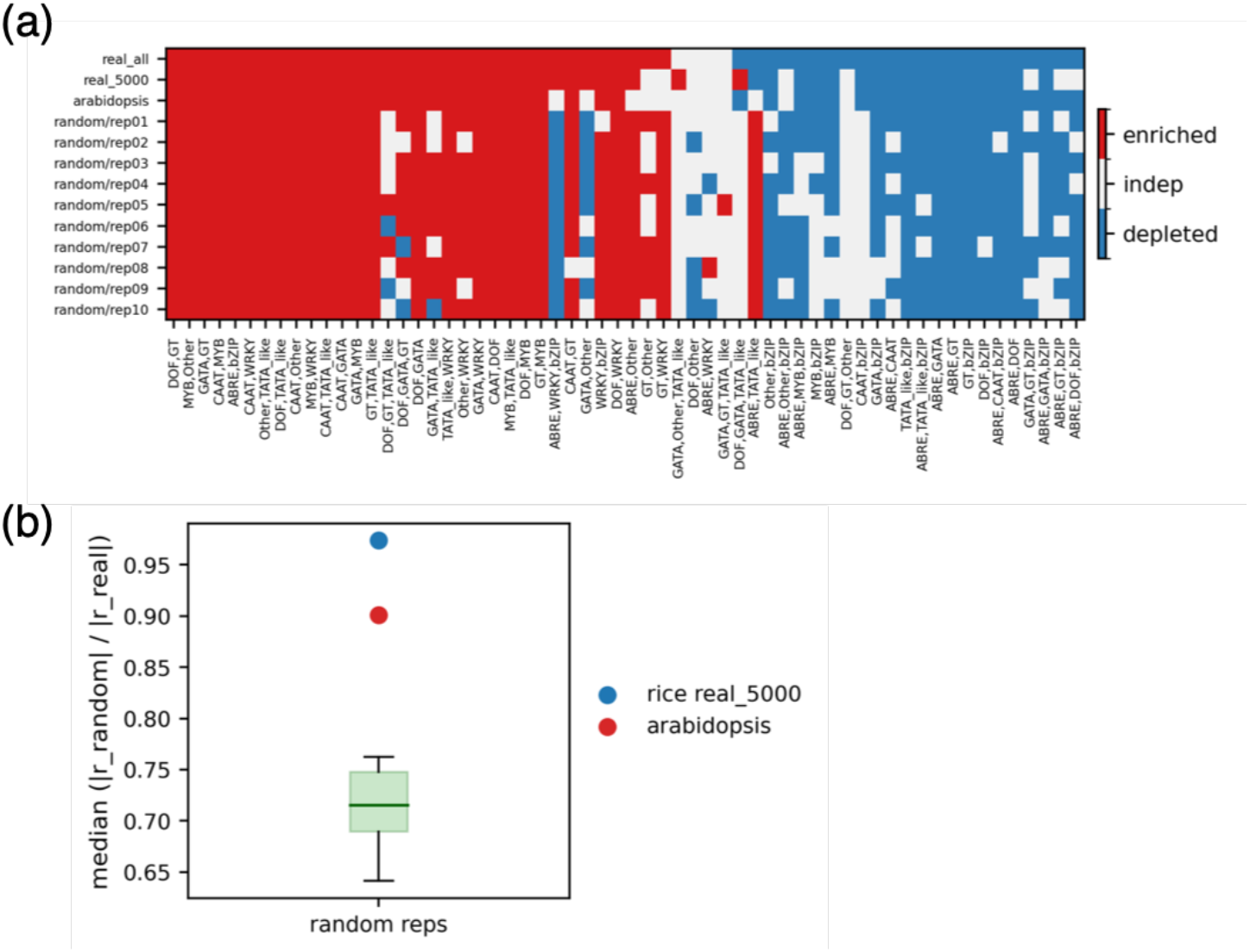
Structured deviations from combinatorial independence within GCIC islands and erosion under randomization. **(a) Dominant combination enrichment and depletion structure**. Sign map of residuals *r* = log_10_ (*f*_observed_/*f*_expected_) for the top-ranked motif-family combinations across real rice, *Arabidopsis*, and randomized replicates, classified using eps = 0.1. Red denotes enrichment relative to independence, blue denotes depletion, and white indicates near-independent behavior. Real genomes exhibit conserved hierarchical combinatorial structure that progressively collapses under randomization. **(b) Quantification of shrinkage toward independence**. Boxplots showing median (|r_random|/|r_real|) across dominant motif-family combinations for randomized replicates, with real rice and *Arabidopsis* values overlaid. Randomization drives systematic erosion of internal combinatorial organization toward independence, whereas real genomes preserve structured deviations.

### Conserved scaling law of combinatorial vocabulary expansion

To quantify the organizational principle underlying GCIC formation, we examined the relationship between total motif-family occurrences per gene and the number of unique motif families represented within GCICs.

In both rice and *Arabidopsis*, vocabulary diversity increased continuously with motif-family abundance, with no evidence of saturation (Fig. 1a). Growth curves were well described by sublinear scaling relations with highly similar exponents across species (b = 0.623 in rice and b = 0.539 in *Arabidopsis*; R^2^ = 0.905 and 0.889, respectively), indicating conserved macroscopic organization of combinatorial landscape expansion.

### Universal hierarchical organization of motif-family combinations

We next analyzed the frequency distribution of motif-family combinations across GCIC islands. In both genomes, combination usage followed strongly skewed hierarchical rank–frequency distributions, with dominant combinations accounting for a large fraction of islands and a long tail of rare combinations persisting (Fig. 1b).

Within the principal rank regime examined, distributions were well approximated by comparable scaling relations across species (α = 3.444 in rice and α = 3.452 in *Arabidopsis*; R^2^ = 0.868 and 0.848, respectively), demonstrating conserved hierarchical organization of combinatorial usage.

### GCICs are predominantly intergenic with broadly similar compartmental distributions across species

To connect GCIC organization with genomic regulatory structure, we quantified the distribution of GCIC islands across promoters, exons, introns, and intergenic regions.

In both rice and *Arabidopsis*, GCICs were broadly distributed outside core promoters, with the largest fraction residing in intergenic regions (Fig. 1c). Despite species-specific differences in gene structure, the overall compartmental composition of GCICs appeared broadly similar across species.

### Conservation of dominant combinatorial identities across species

While macroscopic scaling laws and hierarchical distributions demonstrate conserved statistical organization, we next examined whether the dominant motif-family combinations themselves were preserved.

Comparison of the top-ranked combinations in rice and *Arabidopsis* revealed substantial overlap across a wide range of ranks, with more than 60–90% of dominant combinations shared depending on the cutoff threshold (Fig. 2a), indicating conservation of core combinatorial identities.

### Quantitative correspondence of combination usage frequencies

Frequencies of shared motif-family combinations displayed strong proportional correspondence between rice and *Arabidopsis*, spanning multiple orders of magnitude (Fig. 2b).

Linear regression in log–log space revealed near-proportional conservation of combination deployment across genomes (slope = 0.996, R^2^ = 0.938), demonstrating that dominant combinatorial identities are conserved not only in presence but also in quantitative usage.

### Disruption of higher-order GCIC architectures under randomization

We next examined whether GCIC architectural organization persists under sequence randomization.

Analysis of GCIC multiplicity per gene revealed that real genomes harbor a substantial fraction of genes containing multiple GCIC islands, forming higher-order architectures across a broad dynamic range of GCIC multiplicity (Fig. 3). In contrast, randomized rice subsets exhibited systematic collapse toward single-island configurations, with strong enrichment of GCIC = 1 genes and depletion of multi-island structures.

This collapse extended across the distribution, including the low-order regime (GCIC = 1–4; Fig. 3a) and the heavy-tail regime of higher-order architectures (GCIC ≥ 3; Fig. 3b). Jensen–Shannon divergence analysis further confirmed that randomized distributions were consistently distant from real genomic landscapes, whereas cross-species distributions remained highly similar (Fig. 3c).

### Collapse of motif-family combinatorial structure toward independence under randomization

To determine whether disruption of GCIC architectures mechanistically originates from loss of internal combinatorial organization, we quantified deviations of motif-family combination frequencies from an independence model. In real rice and *Arabidopsis* genomes, dominant motif-family combinations consistently exhibited strong enrichment or depletion relative to independent expectations, forming a conserved hierarchical preference structure (Fig. 4a). In contrast, randomization progressively erased these structured deviations, driving combination frequencies toward independence.

Quantification of deviation magnitude revealed systematic contraction of residual amplitudes under randomization, whereas real subsampled genomes largely preserved the original combinatorial structure. Median shrinkage toward independence, defined as median (|r_random|/|r_real|), was approximately 0.7 across randomized replicates (corresponding to ∼30% contraction), while remaining near unity in real rice and at similarly high levels in *Arabidopsis* (Fig. 4b).

## Discussion

A central result of this study is the cross-genome conservation of macroscopic statistical descriptors of GCIC landscapes under biologically grounded gene-centered observational resolution (Ohmori, 2026). Rather than focusing on individual regulatory elements, we demonstrate that GCIC organization is characterized by reproducible scaling laws, hierarchical rank–frequency distributions, conserved dominant motif-family combinations, and higher-order architectural organization across plant genomes.

In both rice and *Arabidopsis*, combinatorial vocabulary diversity follows highly similar sublinear scaling relations with motif-family abundance (Fig. 1a), quantified by comparable scaling exponents, indicating open-ended yet constrained expansion of regulatory combinatorial space. Likewise, motif-family combination usage exhibits conserved hierarchical rank–frequency organization (Fig. 1b) with near-identical scaling behavior across species. Quantitative correspondence of shared dominant combinations further demonstrates that GCIC organization is conserved not only structurally but also in its deployment across genomes (Fig. 2b).

Together, these macroscopic regularities define a conserved statistical regulatory phase characterized by invariant scaling relations, hierarchical organization, and stable combinatorial identities.

### Emergence and disruption of hierarchical GCIC architecture

While GCIC landscapes exhibit conserved macroscopic organization across species, randomization analyses reveal that this organization depends on native sequence structure.

Real genomes maintain extensive higher-order GCIC architectures composed of multiple islands per gene spanning a broad dynamic range (Fig. 3a–c). In contrast, randomized rice subsets undergo systematic collapse toward single-island configurations, with pronounced depletion of multi-island structures across the full distribution. This architectural collapse mirrors the loss of internal combinatorial organization revealed by progressive contraction of motif-family interaction residuals toward independence, as quantified by shrinkage ratios (Fig. 4a,b).

These results indicate that higher-order GCIC architecture emerges directly from hierarchical motif-family interactions within islands, and that disruption of internal combinatorial structure destabilizes GCIC organization at the genomic scale.

### GCICs as emergent statistical regulatory landscapes

Recent functional and evolutionary studies increasingly challenge the view of gene regulation as being encoded by discrete, individually conserved cis-regulatory elements. Instead, regulatory output often emerges from distributed collections of weak and partially redundant enhancers, conferring robustness despite extensive turnover of individual regulatory sequences (Frankel et al., 2010; Cannavò et al., 2016; Osterwalder et al., 2018; Kvon et al., 2014).

Consistent with this perspective, genome-wide reporter assays demonstrate that regulatory potential is broadly distributed across genomic sequence space rather than confined to annotated elements (Arnold et al., 2013; Muerdter et al., 2015). Enhancer activity further depends on probabilistic combinatorial motif organization rather than deterministic high-affinity binding sites (Spitz & Furlong, 2012; Crocker et al., 2016; Farley et al., 2015; Heinz et al., 2015).

Within this framework, GCICs capture large-scale combinatorial structure generated by motif-family overlap in gene-centered coordinates, representing regulatory landscapes rather than fixed sequence modules. Previous analyses established that GCICs are anchored to functional cis-regulatory space, overlapping extensively with experimentally characterized regulatory regions, conserved noncoding sequences, and transcriptionally active regulatory elements (Ohmori, 2026).

### A fluctuation-structured regulatory code grounded in sequence statistics

The conserved statistical phase observed across divergent plant genomes indicates an organizational principle rooted in intrinsic sequence-level fluctuations. Intrinsic sequence statistics naturally generate motif clustering and combinatorial regulatory potential (Stormo & Fields, 1998; Segal et al., 2008; Wunderlich & Mirny, 2009; Estrada et al., 2016), from which stable macroscopic regulatory landscapes emerge when viewed in motif-family space.

Randomization experiments demonstrate that disruption of native sequence correlations drives combinatorial organization toward independence, indicating that hierarchical motif-family interactions are maintained by intrinsic sequence structure rather than unconstrained motif accumulation (Stormo & Fields, 1998; Segal et al., 2008; Wunderlich & Mirny, 2009; Estrada et al., 2016).

This erosion of structured interactions under randomization highlights the probabilistic nature of combinatorial regulatory logic (Spitz & Furlong, 2012; Crocker et al., 2016; Farley et al., 2015; Heinz et al., 2015).

Together, these findings support regulatory systems operating within probabilistic sequence spaces rather than deterministic motif codes (Levine & Davidson, 2005; Crocker et al., 2016).

### Statistical origin of regulatory robustness within GCICs

A central puzzle in regulatory evolution is how gene expression programs remain stable despite extensive enhancer turnover (Wittkopp & Kalay, 2011; Villar et al., 2015). Robustness increasingly appears to arise from integration of multiple weak and partially redundant regulatory inputs rather than reliance on single dominant cis-elements (Frankel et al., 2010; Cannavò et al., 2016; Osterwalder et al., 2018).

Within GCIC landscapes, robustness emerges from combinatorial redundancy, where overlapping motif families collectively generate probabilistic regulatory potential. Loss of internal combinatorial structure destabilizes higher-order GCIC architectures, underscoring the role of hierarchical motif-family interactions in maintaining regulatory landscape integrity.

### Probabilistic generation, opportunistic engagement, and selective stabilization

GCIC landscapes likely arise probabilistically as natural consequences of sequence statistics, without requiring functional selection at their origin. Transcription factor binding and chromatin accessibility can opportunistically engage pre-existing combinatorial landscapes, enabling regulatory activity where sufficient motif-family overlap exists.

When particular configurations confer advantageous effects, selection may stabilize these regions as conserved cis-regulatory elements, while most GCIC landscapes remain fluctuating across evolutionary time.

### Distinction from unconstrained motif clustering

Unstructured motif accumulation would be expected to produce saturation of motif-family diversity and uniform combinatorial usage. In contrast, GCIC landscapes exhibit continuous vocabulary expansion, hierarchical rank–frequency distributions, conserved dominant combinations, and characteristic collapse of organization under randomization, indicating a structured organizational regime rather than unconstrained randomness.

### GCICs as a conserved statistical regulatory phase

In statistical physics, phases are defined by invariant macroscopic properties emerging from collective microscopic interactions. GCIC landscapes exhibit conserved scaling relations, hierarchical combinatorial organization, stable dominant interactions, and reproducible responses to perturbation across genomes, consistent with phase-like behavior in regulatory sequence space.

Evolution thus appears to conserve macroscopic statistical organization of regulatory landscapes rather than specific regulatory sequences, with functional elements representing stabilized realizations of this probabilistic phase.

### Toward a statistical theory of regulatory landscapes

Epigenomic and functional mapping has revealed expansive and dynamic regulatory chromatin landscapes across animals and plants (Roadmap Epigenomics Consortium, 2015; Moore et al., 2026; Oka et al., 2017; Sun et al., 2019; Ricci et al., 2019). GCICs provide a sequence-based statistical framework explaining how such landscapes emerge naturally while remaining evolutionarily flexible.

By shifting regulatory conservation from individual elements to macroscopic statistical organization, the fluctuation-structured regulatory code integrates enhancer turnover, combinatorial redundancy, and robustness into a unified theoretical framework.

## Acknowledgements

This work was supported by the Cabinet Office, Government of Japan, Moonshot Research and Development Program for Agriculture, Forestry and Fisheries (funding agency: Bio-oriented Technology Research Advancement Institution), Grant Number JPJ009237.

## References

Arnold, C.D., Gerlach, D., Stelzer, C., Boryń, Ł.M., Rath, M. & Stark, A. Genome-wide quantitative enhancer activity maps identified by STARR-seq. Science 339, 1074–1077 (2013).

Berardini, T.Z. et al. The Arabidopsis Information Resource: making and mining the “gold standard” annotated reference plant genome. Genesis 53, 474–485 (2015).

Cannavò E, Khoueiry P, Garfield DA, Geeleher P, Zichner T, Gustafson EH, Ciglar L, Korbel JO, Furlong EE. Shadow enhancers are pervasive features of developmental regulatory networks. Curr. Biol. 26, 38–51 (2016).

Crocker, J., Noon, E.P. & Stern, D.L. The soft touch: low-affinity transcription factor binding sites in development and evolution. Trends Genet. 32, 581–591 (2016).

Deyholos, M.K. & Sieburth, L.E. Separable whorl-specific expression and negative regulation by enhancer elements within the AGAMOUS second intron. Plant Cell 12, 1799–1810 (2000).

Estrada, J., Wong, F., DePace, A. & Gunawardena, J. Information integration and energy expenditure in gene regulation. Cell 166, 234–244 (2016).

Farley EK, Olson KM, Zhang W, Brandt AJ, Rokhsar DS, Levine MS. Suboptimization of developmental enhancers. Science 350, 325–328 (2015).

Frankel, N., Davis, G.K., Vargas, D., Wang, S., Payre, F. & Stern, D.L. Phenotypic robustness conferred by apparently redundant transcriptional enhancers. Nature 466, 490–493 (2010).

Heinz, S., Romanoski, C.E., Benner, C. & Glass, C.K. The selection and function of cell type–specific enhancers. Nat. Rev. Mol. Cell Biol. 16, 144–154 (2015).

Howe, K.L. et al. Ensembl Genomes 2020—enabling non-vertebrate genomic research. Nucleic Acids Res. 48, D689–D695 (2020).

Kawahara, Y. et al. Improvement of the Oryza sativa Nipponbare reference genome using next generation sequence and optical map data. Rice 6, 4 (2013).

Kvon EZ, Kazmar T, Stampfel G, Yáñez-Cuna JO, Pagani M, Schernhuber K, Dickson BJ, Stark A. Genome-scale functional characterization of Drosophila developmental enhancers in vivo. Nature 512, 91–95 (2014).

Levine, M. & Davidson, E.H. Gene regulatory networks for development. Proc. Natl Acad. Sci. USA 102, 4936–4942 (2005).

Lin, J. Divergence measures based on the Shannon entropy. IEEE Trans. Inf. Theory 37, 145–151 (1991).

Melnikov, A., Murugan, A., Zhang, X. et al. Systematic dissection and optimization of inducible enhancers in human cells using a massively parallel reporter assay. Nat. Biotechnol. 30, 271–277 (2012).

Moore, J.E., Pratt, H.E., Fan, K., Phalke, N., Fisher, J., Elhajjajy, S.I., Andrews, G., Gao, M., Shedd, N., Fu, Y., Lacadie, M.C., Meza, J., Ganna, M., Choudhury, E., Swofford, R., Farrell, N.P., Pampari, A., Ramalingam, V., Reese, F., Borsari, B., Yu, M., Wattenberg, E., Ruiz-Romero, M., Razavi-Mohseni, M., Xu, J., Galeev, T., Beer, M.A., Guigó, R., Gerstein, M.B., Engreitz, J.M., Ljungman, M., Reddy, T.E., Snyder, M.P., Epstein, C.B., Gaskell, E., Bernstein, B.E., Dickel, D.E., Visel, A., Pennacchio, L.A., Mortazavi, A., Kundaje, A. & Weng, Z. An expanded registry of candidate cis-regulatory elements for studying transcriptional regulation. Nature (2026). 10.1038/s41586-025-09909-9

Muerdter, F., Boryń, Ł.M. & Arnold, C.D. STARR-seq—principles and applications. Genomics 106, 145–150 (2015).

Ohmori, Y., Toriba, T., Nakamura, H., Ichikawa, H. & Hirano, H.-Y. Temporal and spatial regulation of DROOPING LEAF gene expression that promotes midrib formation in rice. Plant J. 65, 77–86 (2011).

Ohmori, Y. Gene-centered identification of cis-regulatory islands reveals regulatory landscapes complementary to motif-centric approaches. Preprint at 10.64898/2026.01.05.697455 (2026).

Oka, R., Zicola, J., Weber, B., Anderson, S.N., Hodgman, C., Gent, J.I., Wesselink, J.J., Springer, N.M., Hoefsloot, H.C.J., Turck, F. & Liu, Z. Genome-wide mapping of transcriptional enhancer activity in rice using STARR-seq. Genome Biol. 18:137 (2017).

Osterwalder M, Barozzi I, Tissières V, Fukuda-Yuzawa Y, Mannion BJ, Afzal SY, Lee EA, Zhu Y, Plajzer-Frick I, Pickle CS, Kato M, Garvin TH, Pham QT, Harrington AN, Akiyama JA, Afzal V, Lopez-Rios J, Dickel DE, Visel A, Pennacchio LA. Enhancer redundancy provides phenotypic robustness in mammalian development. Nature 554, 239–243 (2018).

Ricci, W.A., Lu, Z., Ji, L., Marand, A.P., Ethridge, C.L., Murphy, N.G., Noshay, J.M., Galli, M., Mejía-Guerra, M.K., Colomé-Tatché, M., Schmitz, R.J., Zhai, J., Scanlon, M.J., Buckler, E.S., Springer, N.M. & Ross-Ibarra, J. Widespread long-range cis-regulatory elements in the maize genome. Nat. Plants 5, 1237–1249 (2019).

Roadmap Epigenomics Consortium. Integrative analysis of 111 reference human epigenomes. Nature 518, 317–330 (2015).

Segal, E., Raveh-Sadka, T., Schroeder, M., Unnerstall, U. & Gaul, U. Predicting expression patterns from regulatory sequence in Drosophila segmentation. Nature 451, 535–540 (2008).

Spitz, F. & Furlong, E.E.M. Transcription factors: from enhancer binding to developmental control. Nat. Rev. Genet. 13, 613–626 (2012).

Stormo, G.D. & Fields, D.S. Specificity, free energy and information content in protein– DNA interactions. Trends Biochem. Sci. 23, 109–113 (1998).

Sun, J., He, N., Niu, L., Huang, Y., Shen, W., Zhang, Y., Li, L., Hou, C., Wang, X., Zhang, H., Chen, Z.J. & Zhu, J.K. Global Quantitative Mapping of Enhancers in Rice by STARR-seq. Genomics Proteomics Bioinformatics 17, 140–153 (2019).

Villar D, Berthelot C, Aldridge S, Rayner TF, Lukk M, Pignatelli M, Park TJ, Deaville R, Erichsen JT, Jasinska AJ, Turner JM, Bertelsen MF, Murchison EP, Flicek P, Odom DT. Enhancer evolution across 20 mammalian species. Cell 160, 554–566 (2015).

Wittkopp, P.J. & Kalay, G. Cis-regulatory elements: molecular mechanisms and evolutionary processes underlying divergence. Nat. Rev. Genet. 13, 59–69 (2011).

Wunderlich, Z. & Mirny, L.A. Different gene regulation strategies revealed by analysis of binding motifs. Trends Genet. 25, 434–440 (2009).

